# Prior expectations of motion direction modulate early sensory processing

**DOI:** 10.1101/2020.03.06.980672

**Authors:** Fraser Aitken, Georgia Turner, Peter Kok

## Abstract

Perception is a process of inference, integrating sensory inputs with prior expectations. However, little is known regarding the temporal dynamics of this integration. It has been proposed that expectation plays a role early in the perceptual process, by biasing early sensory processing. Alternatively, others suggest that expectations are integrated only at later, post-perceptual decision-making stages. The current study aimed to dissociate between these hypotheses. We exposed male and female human participants (N=24) to auditory cues predicting the likely direction of upcoming noisy moving dot patterns, while recording millisecond-resolved neural activity using magnetoencephalography (MEG). First, we found that participants’ reports of the moving dot directions were biased towards the direction predicted by the auditory cues. To investigate when expectations affected sensory representations, we used inverted encoding models to decode the direction represented in early sensory signals. Strikingly, the auditory cues modulated the direction represented in the MEG signal as early as 150ms after visual stimulus onset. This early neural modulation was related to perceptual effects of expectation: participants with a stronger perceptual bias towards the predicted direction also revealed a stronger reflection of the predicted direction in the MEG signal. For participants with this perceptual bias, a trial-by-trial correlation between decoded and perceived direction already emerged prior to visual stimulus onset (∼-150ms), suggesting that the pre-stimulus state of the visual cortex influences sensory processing. Together, these results suggest that prior expectations can influence perception by biasing early sensory processing, making expectation a fundamental component of the neural computations underlying perception.

**Significance statement:** Perception can be thought of as an inferential process in which our brains integrate sensory inputs with prior expectations to make sense of the world. This study investigated whether this integration occurs early or late in the process of perception. We exposed human participants to auditory cues which predicted the likely direction of visual moving dots, while recording neural activity with millisecond resolution using magnetoencephalography (MEG). Participants’ perceptual reports of the direction of the moving dots were biased towards the predicted direction. Additionally, the predicted direction modulated the neural representation of the moving dots just 150 ms after they appeared. This suggests that prior expectations affected sensory processing at very early stages, playing an integral role in the perceptual process.

## Introduction

Since Helmholtz (1867) described perception as a process of ‘unconscious inference’, it has become widespread to conceptualise perception as an integration of bottom-up sensory information with top-down prior expectations (Summerfield and De Lange, 2014; Kersten and Yuille, 2003; Friston, 2005). However, the neural mechanisms and time-course of this integration remain controversial. On the one hand, influential theories of predictive processing, such as predictive coding (Rao and Ballard, 1999; Friston, 2005), posit that top-down predictions are integrated with bottom-up sensory information from the moment inputs arrive in the cortex, such that sensory representations are modulated by expectations at early sensory stages (Rao and Ballard, 1999; Friston, 2005; Lee and Mumford, 2003; Wyart et al., 2012; Keller and Mrsic-Flogel, 2018). In support of this hypothesis, many studies have shown that prior expectations can modulate processing at the earliest stages of the cortical hierarchy (Kok et al., 2012, Alink et al., 2010, Den Ouden et al., 2009), as well as early in time (starting around 100-150 ms post-stimulus; Todorovic et al., 2011, Alilović et al., 2019, Wacongne et al., 2011, Hsu et al., 2015, Jabar et al., 2017, Aru et al., 2016), even prior to stimulus presentation (Kok et al., 2017; Sherman et al., 2016).

Alternatively, it has been suggested that prior expectations leave early sensory processing untouched, and instead only modulate later decision-making processes (Rungratsameetaweemana et al., 2018; Rao et al.,2012; Bang and Rahnev, 2017), for instance in parieto-frontal brain circuits (Gold and Shadlen, 2007; Heekeren et al., 2004). Under this account, effects of expectations in early sensory regions as revealed by previous functional magnetic resonance imaging (fMRI) studies are proposed to reflect late, post-decision feedback signals, simply ‘informing’ sensory regions of the decision that has been made. Even early effects of expectations revealed by electrophysiological studies (Todorovic et al., 2011; Chaumon et al., 2008; Gamond et al., 2011) may be epiphenomena rather than directly impacting perception, analogous to the proposals that working memory representations in sensory regions reflect epiphenomena (Xu, 2018; but see Zhang et al., 2019).

Previous studies have been unable to distinguish between these two hypotheses, since they have not linked neural effects of expectation to behavioural changes in perception. Additionally, most previous electrophysiological studies have studied changes in the overall amplitude of the neural response to (un)expected stimuli, rather than probing stimulus-specific representations in the neural signal (Rungratsameetaweemana et al., 2018; Aru et al., 2016; Alilović et al., 2019). This is critical, since previous studies have shown that informational content can be fully dissociated from the overall amplitude of neural signals (Kok et al., 2012, Harrison and Tong, 2009). Therefore, these studies may have missed stimulus-specific effects of expectations on sensory processing.

Here, we overcame these limitations by using magnetoencephalography (MEG) to directly relate effects of expectation on neural representations to effects on the contents of perception. Participants were exposed to auditory cues which, unbeknownst to them, predicted the likely motion direction of a subsequent random dot kinetogram (RDK). Perception was probed by asking participants to report which direction the dots were moving in. A forward model decoder (Brouwer and Heeger, 2009, Kok et al., 2017), trained on task-irrelevant RDKs presented in independent runs, was used to reveal the motion direction represented in the MEG signal immediately after stimulus presentation. This allowed us to determine at which time-points the sensory representation was modulated by the prediction cue.

To preview, we found that prior expectations modulated the content of sensory presentations as early as 150 ms post-stimulus. These neural effects were mirrored by a bias in perception, in line with proposals that expectations can bias perception by modulating early sensory processing.

## Materials and methods

### Participants

30 healthy human volunteers participated in the MEG experiment. The study was approved by the UCL Research Ethics Committee, and all participants gave informed consent and received monetary compensation. Two participants were excluded because of excessive head movement, one for excessive eye blink artefacts during stimulus presentation, and three due to below-threshold task performance (*r* < 0.9 between mean perceived and presented direction). The remaining 24 participants (11 female; age 25 ± 8; mean ± SD) had normal or corrected-to-normal vision. This sample size was chosen on the basis of similar previous studies which had observed significant effects (Kok et al. 2013; Kok et al., 2017; Mostert et al., 2015).

### Stimuli

All stimuli were generated using MATLAB (MathWorks, Natick, MA, USA, RRID:SCR_001622) and the Psychophysics Toolbox (Brainard, 1997, RRID:SCR_002881). The visual stimuli were RDKs, which consisted of white dots (0.1° visual angle dot size, 2.5 dots per square degree) on a grey background. Each RDK display contained a given proportion of dots moving in a coherent direction, with the remaining dots moving in random directions. Each dot appeared at a random location, moved at a speed of 6°/s, and lasted for 200ms before disappearing. The dots were displayed in an annulus (inner diameter, 3°; outer diameter, 15°), surrounding a white fixation bullseye (diameter: 0.7°) for 1s. The auditory stimuli consisted of pure tones (450 or 1000 Hz) and lasted 200ms.

During the behavioural session, visual stimuli were presented on an LCD monitor (1024 × 768 resolution, 60 Hz refresh rate) and tones were presented on external speakers. During the MEG session, visual stimuli were projected on a screen placed 58 cm from the participants’ eyes (1024 × 768 resolution, 60 Hz refresh rate), and auditory stimuli were presented via earphones inserted into the ear canal (E-A-RTONE 3A 10 Ω, Etymotic Research).

### Experimental procedure

The experiment consisted of two types of task runs. In the main task, each trial consisted of an auditory cue (200 ms) followed after 550 ms by a visual RDK stimulus for 1000 ms (Figure 1A). After a 500 ms interval, participants reported the direction of the coherently moving dots by orienting a line segment in a 360° circle (2500 ms). The initial direction of the line was randomised between −45° and 135°. After the response interval, during the intertrial interval (ITI; 1500 ms), the fixation bullseye was replaced by a single dot, signalling the end of the trial while still requiring participants to fixate. The RDKs had one of five possible directions of coherent motion: 9°, 27°, 45°, 63° or 81°. Participants were informed the coherent direction would range between 0° and 90° but not that there was a discrete set of possible directions. The two auditory cues predicted either 27° or 63°, respectively, with 60% probability (Figure 1B). Participants were not informed of this cue-direction relationship. The four non-predicted directions were each equally likely to occur (10% probability). The relationship between which tone predicted which direction was counterbalanced across participants. Each run contained 60 trials (∼6 min).

**Figure 1.**
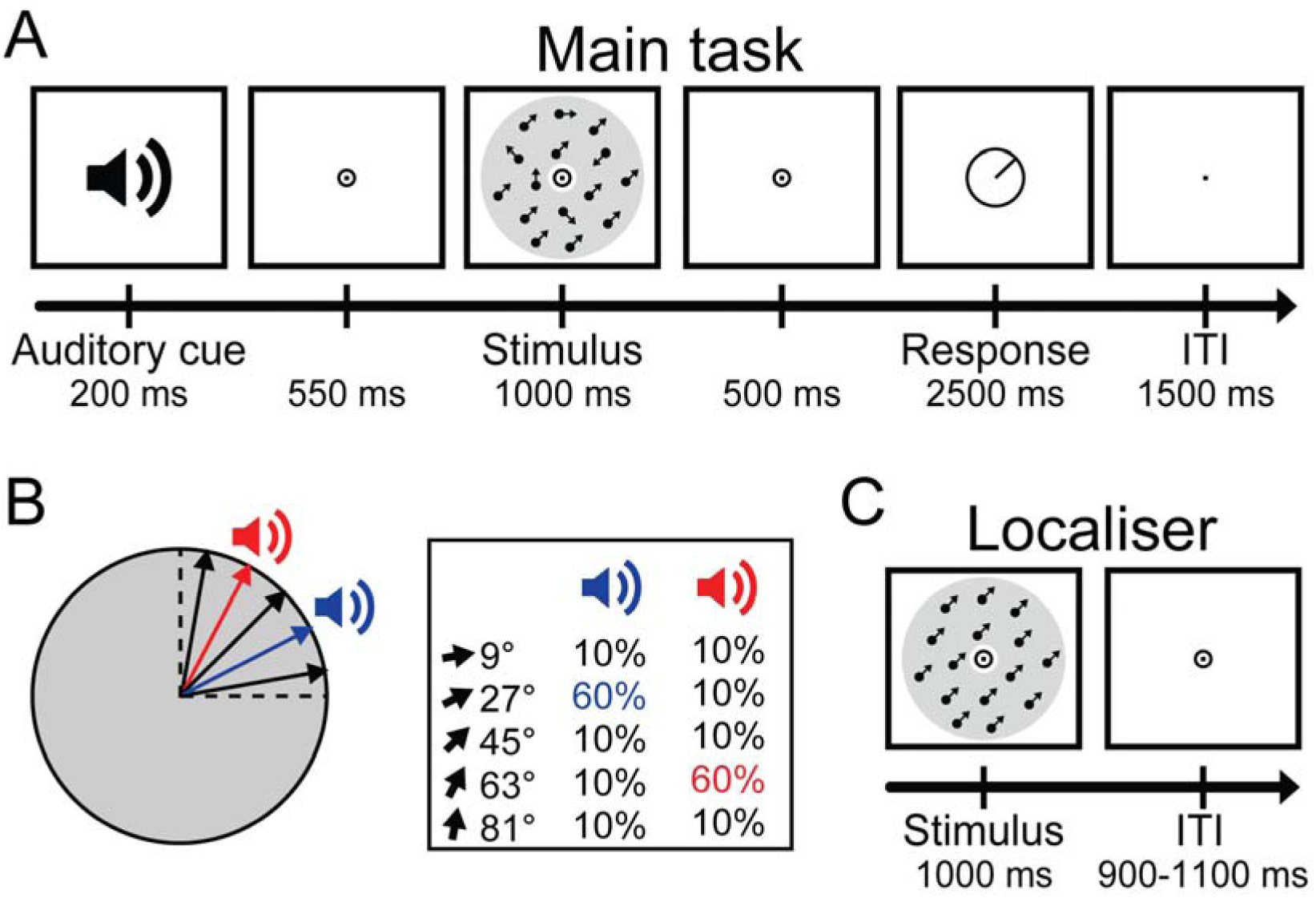
Schematic diagram of the experimental procedure. **(A)** The main task. Participants were presented with an auditory cue, followed by a random dot kinematogram (RDK) stimulus. Participants indicated their response on a continuous scale by rotating a line segment. ITI = Intertrial Interval. **(B)** Possible coherent directions in the RDK. One tone predicted 27°, and the other tone predicted 63°, each with 60% validity. **(C)** In localiser runs, participants were presented with task-irrelevant moving dots stimuli with 100% coherence, while performing a dot dimming task at fixation.

During localiser runs, RDKs were presented with 100% coherence, creating training data for the MEG decoder (Figure 1C). 11 motion directions were presented in a pseudorandom order, for 1000 ms each. These directions were –45°, −27°, −9°, 9°, 27°, 45°, 63°, 81°, 99°, 117°, and 135°. One localiser block consisted of 88 trials (∼3 min). The fixation bullseye at the centre of the annulus dimmed at random time-points, and subjects were instructed to press a button when this occurred. The ITI was jittered between 900 and 1100 ms. During these runs, the moving dots were fully task-irrelevant, in order to extract motion direction signals independent of task demands (Kok et al., 2017). The task was also intended to encourage central fixation, in order to minimise eye-movement related confounds (Mostert et al., 2018).

All participants took part in a behavioural session 1-4 days prior to the MEG session, in order to familiarise them with the task and expose them to the cue-direction contingencies. Participants received written instructions and performed two short blocks (of 20 and 40 trials, respectively) with trial-by-trial feedback to facilitate learning. They then performed 7 main task blocks of 60 trials each (∼45 min) during which they no longer received trial-by-trial feedback, but were informed of their mean error after each block for motivation, as in the MEG session. The RDKs began with 40% coherence in the instructions and practice blocks, to facilitate learning of the task, and gradually reduced from 40% to 20% coherence during the main behavioural session. Finally, participants participated in one localiser block to familiarise them with the fixation dimming task.

In the MEG session, participants performed 5-7 runs (∼9 min each). Each run consisted of 60 trials of the main task, followed by a 15 s pause, then one block of the localiser task. In the main task, the RDKs had 20% coherence. After the experiment, participants filled out a debriefing questionnaire, to verify the implicit nature of the expectations. They were asked: ‘Did any directions of motion occur more often than the rest? If so, please indicate which direction(s) you thought occurred more often than the others.’ Subsequently, participants were asked, ‘Did you notice any relationship between the tones you heard and the directions of motion you saw? If so, please describe the relationship you observed in the text box below.’ For both questions, they were also provided with a unit circle in which they could illustrate their answer by drawing arrows to represent specific motion directions.

### MEG recording and preprocessing

Whole-head magnetic signals were recorded continuously (600 Hz sampling rate) using a CTF MEG system with 272 functioning axial gradiometers inside a magnetically shielded room. Participants were seated upright, and indicated their responses on an MEG-compatible button box. To minimise eye blink-related artefacts, participants were instructed to blink only when the RDK was not on the screen. Eye-movement was recorded using an Eyelink 1000 eye tracker (1000 Hz sampling rate). Presentation latencies for stimuli (visual ∼17 ms; auditory ∼15 ms) were measured using a photodiode and microphone; these were used to align the MEG and eye-tracking data to the onset of stimulus presentation. After the first MEG run, participants were informed of their head motion and encouraged to stay as still as possible during the recordings. Since participants displayed substantially more head motion during the first run, this run was discarded for all participants.

The data were preprocessed using FieldTrip (Oostenveld et al., 2011). To detect irregular artefacts, the variance, collapsed over channels and time, was calculated for each trial. Trials with large variances were visually inspected and removed if they contained large and irregular artefacts. Independent component analysis was used to remove regular artefacts, by correlating the components with the eye-tracking data to identify eye blinks, and then manually inspecting before removing components related to eye blinks. Trials with eye blinks during RDK presentation were removed. Data were low-pass filtered with a two-pass Butterworth filter with a filter order of 6 and a cut off of 40 Hz. The data were baseline-corrected on the interval of −250 ms to 0 ms relative to auditory cue onset for the main task, and −200 ms to 0 ms relative to visual onset for the localiser task.

### Decoding analysis

To probe the effect of expectations on stimulus representations in visual cortex, we used a forward modelling approach (Brouwer and Heeger, 2009) to decode motion directions from the MEG signal (Kok et al., 2017; Myers et al., 2015). This approach has been highly successful at decoding continuous stimulus features from neural data (Brouwer and Heeger, 2009; 2011; Garcia et al., 2013; Myers et al., 2015; Kok et al., 2013; 2017). Furthermore, it yields decoded features on a continuous dimension rather than a discrete classification, making it potentially more sensitive to subtle biases than a categorical classifier.

The decoding approach consisted of two stages. First, the model was trained on the MEG data from the moving dots localiser to create an encoding model: a transformation from stimulus (motion direction) space to MEG sensor space. Then, this encoding model was inverted to create a decoding model, which was used to transform unseen MEG data (from the main task runs) from sensor space to motion direction space. Thus, the decoding model was estimated on the basis of the moving dot localiser data, and then applied to the data from the main experiment to generalise from sensory signals evoked by task-irrelevant moving dots to the noisy moving dot signals evoked in the main task (Kok et al., 2017). To test the performance of the model, we also applied it to the localiser data themselves using a cross-validation approach: in each iteration, one run of the localiser was used as the test set, and the remaining data were used as the training set.

The forward encoding model consisted of 21 hypothetical channels, each with an idealised direction tuning curve: a half-wave rectified sinusoid raised to the sixth power. The 21 channels were spaced evenly within the 180° space ranging from −45° to 135°, to cover all directions presented in the localiser runs. For each participant, the MEG data from the localiser were used to calculate an encoding model. First, a matrix **C**_*train*_ (21 channels x *n*_*train*_ trials) was generated, containing the hypothesised channel amplitudes for each trial. Specifically, each row of matrix **C**_*train*_ was calculated by expressing the presented direction as a hypothetical amplitude for each channel, resulting in the row vector **c**_*train,i*_ of length *n*_*train*_ for each channel *i*. The sensor data were represented in a matrix **B**_*train*_ (272 sensors x *n*_*train*_ trials). The key aspect of the encoding model was a weight matrix, specifying the transformation from stimulus space (represented in matrix **C**_*train*_) to sensor space (matrix B_*train*_). The rows of the weight matrix were calculated by least squares estimation for each channel:

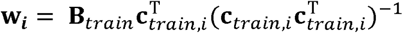

This was used to create a linear encoding model:

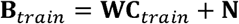

Here, **W** is a weight matrix (272 sensors x 21 channels) specifying the transformation from stimulus representational space (channel activities) to neural representational space (sensor amplitudes). **N** represents the residuals.

In the second stage of the analysis, the decoding model was created by inverting the encoding model. This was achieved using a recently developed method taking the noise covariance between (neighbouring) sensors into account, which increases decoding accuracy compared to a decoding model that does not adjust for noise covariance (Kok et al., 2017; Mostert et al., 2015). First, **B**_***traln***_ and **C**_***traln***_ were demeaned, so that their average over trials was 0 for all sensors and channels, respectively (Kok et al., 2017). Then, noise covariance, *Σ*_i_, between the sensors was estimated for each channel *i* using the following equation:

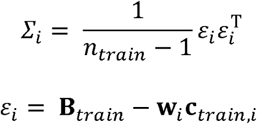

To optimise noise suppression, regularisation by shrinkage, using analytically determined optimal shrinkage parameters, was used to calculate regularised covariance matrices for each channel, 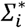 (details are in Blankertz et al., 2011). These regularised covariance matrices were used to create spatial filters. The optimal spatial filter **v**_*i*_ for the *i*^*th*^ channel was estimated as follows (Mostert et al. 2015; Kok et al., 2017):

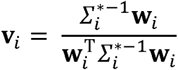

Each filter was normalised so that the magnitude of its output matched the magnitude of the channel activity it would be used to recover. The filters were combined into a decoding weight matrix **V** (272 sensors x 21 channels). This decoding weight matrix could then be used to estimate the channel responses for independent MEG data:

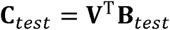

Where **B**_*test*_ (272 sensors x *n*_*test*_ trials) represents the independent test data.

These channel responses were estimated at each time-point of the test data in steps of 5 ms, with the data being averaged within a window of 28.3 ms at each step. The length of 28.3 ms was based on an a priori window of 30 ms, subtracting one sample such that the window contained an odd number of samples and could be centred symmetrically.

To verify the ability to decode motion direction from the MEG signal, we first applied the decoding approach to the localiser data themselves in a ‘leave-one-run-out’ cross-validation method. In each iteration, all localiser runs but one were used to estimate the decoding model, which was then applied to estimate the channel responses in the remaining run. The estimated channel responses were used to compute a weighted average of the 21 basis functions, and the direction at which the resulting curve reached its maximum value constituted the decoded motion direction. Decoding performance was quantified, per time step, by calculating the Pearson correlation between the 11 presented directions, and the mean decoded direction per presented direction. This was repeated with each iteration leaving a different localiser run out, and final decoding performance was quantified by averaging results across all these iterations. We determined the earliest time-point at which decoding performance was significantly above chance at the group level (∼100 ms; see Figure 3), and determined each participant’s individual decoding peak within 10 ms of this group peak (90 – 110 ms post-stimulus). For each participant, the final decoding model was trained on this individual peak time-point (plus the two neighbouring time-points on either side, for robustness) in the localiser data, to optimise detection of early sensory signals. This decoding model was applied to the data from the main task in steps of 5 ms, with the data being averaged within a window of 28.3 ms at each step. As before, the decoded motion direction was calculated as the peak of the curve generated by taking a weighted average of the basis functions, with the estimated channel responses constituting the weights. This procedure yielded a 2D matrix (time x *n*_*test*_) specifying the estimated motion direction for each trial in the main experiment, in a time-resolved manner.

### Statistical analysis

To quantify overall behavioural performance, the mean of all reported directions per presented direction was calculated for each participant (Figure 2A), and used to calculate the Pearson’s correlation coefficient between reported and presented direction, per participant. Participants with a coefficient < 0.9 (N=3) were excluded from further analysis.

**Figure 2.**
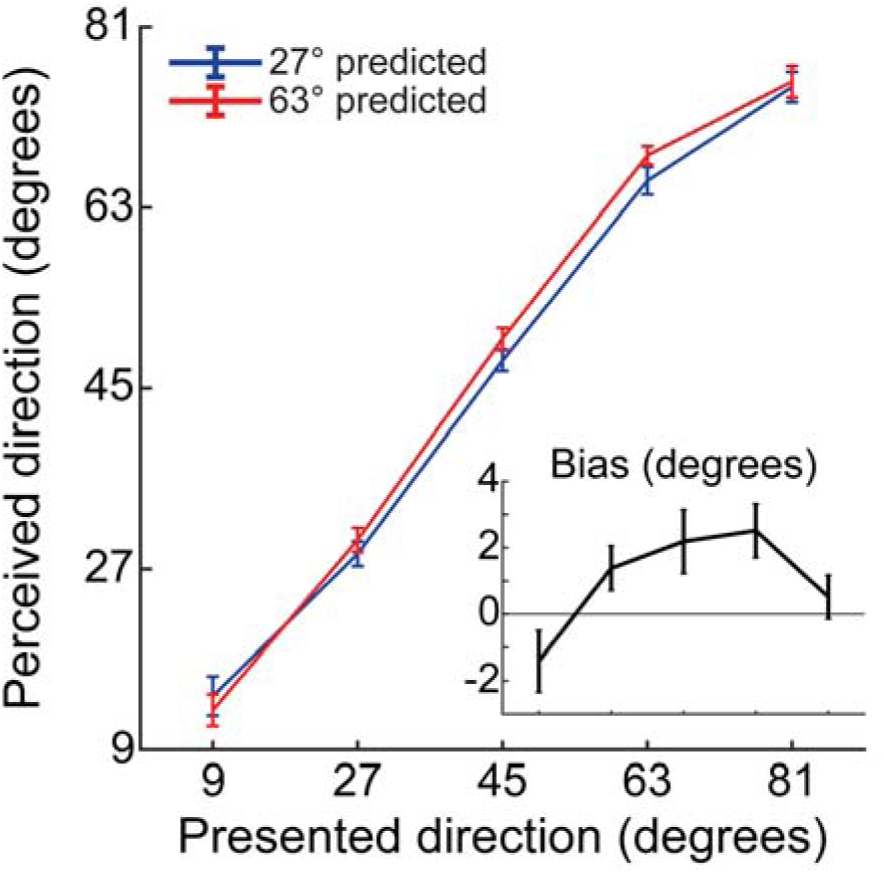
Behavioural results. Reported direction as a function of presented direction. Mean reported direction plotted against presented direction, separately for the two predictive cues. Inset shows the difference between the two cue conditions, i.e., the bias induced by expectation. Error bars indicate standard error of the mean (SEM).

**Figure 3.**
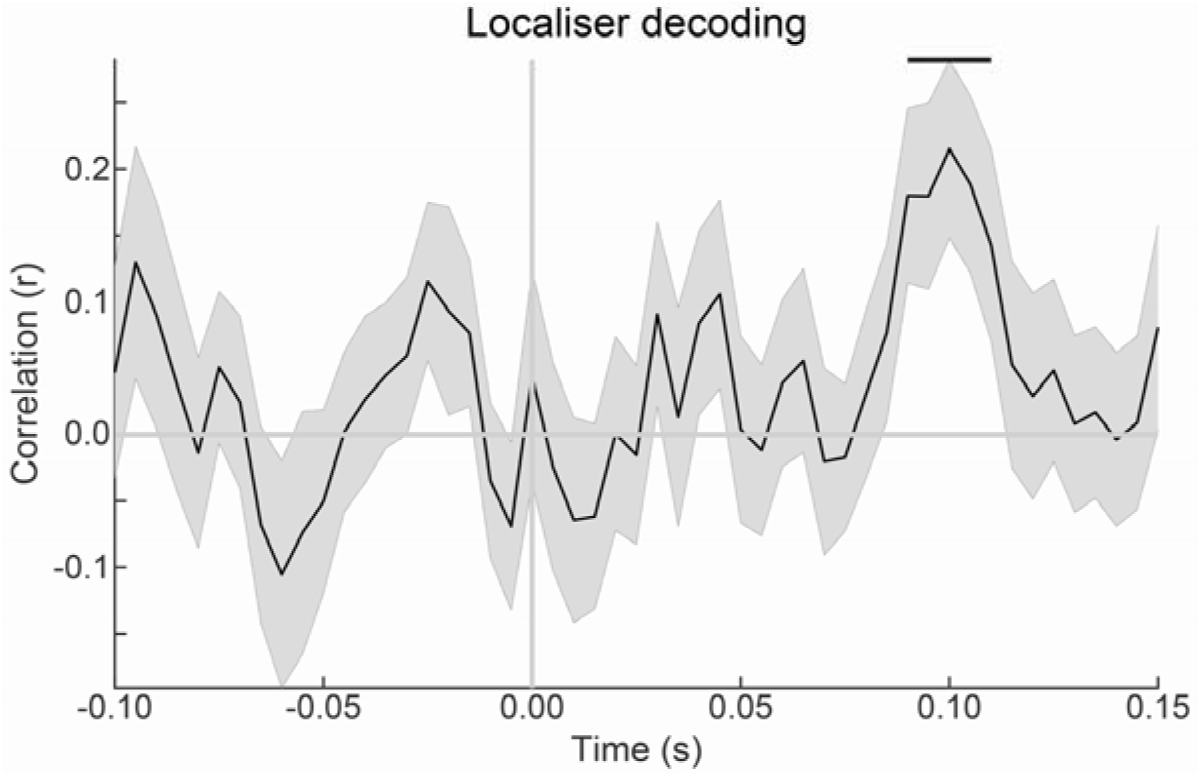
Within-localiser MEG decoding performance. Correlation between presented and decoded direction (cross-validated for each participant, then averaged across participants), plotted for each time-point. The shaded region indicates SEM; horizontal lines represent significant clusters (*p* < 0.05).

To examine the effects of the predictive cues on reported direction, we performed a two-way repeated-measures ANOVA with factors ‘Presented Direction’ and ‘Predicted Direction’. Significant main effects and interactions were followed up with t-tests.

Our main research question was whether prior expectations modulated the neural representation of the moving dot stimuli, and if so, at which time-points. To address this question, we first averaged decoding results over trials, per presented and predicted direction, and then performed a linear subtraction of the decoded direction in conditions where 27° was cued, from the decoded direction in conditions where 63° was cued. The logic behind this method was that, as the only difference between the conditions was the predicted direction, the subtraction would subtract out any signals in common between the cued conditions, and isolate any difference related to the cues. We used cluster-based permutation tests (Maris and Oostenveld, 2007) to establish at which time-points this subtraction was significantly different from zero. Specifically, univariate *t* statistics were calculated for time-points from −250 ms to 500 ms relative to moving dots onset, in 5 ms steps, and neighbouring elements that passed a threshold value corresponding to a *p* value of 0.05 (one tailed) were collected into clusters. Cluster-level test statistics consisted of the sum of t values within each cluster, which were compared to a null distribution created by drawing 10,000 random permutations of the observed data. A cluster was considered significant when its *p* value was below 0.05 (i.e., a cluster of its size occurred in fewer than 5% of the null distribution clusters).

To investigate whether the motion direction signals we decoded were directly related to subjective perception, we calculated the partial correlation between the decoded direction from the MEG data and the perceived direction in each trial, regressing out the presented direction. This analysis was performed separately for participants with a positive behavioural bias induced by the expectation cues (N=17) and those without such a bias (N=7). Statistical tests of correlation coefficients were preceded by applying Fisher’s *r*-to-Z transform (Fisher, 1915), and the resulting time-courses of Z values were subjected to cluster-based permutation tests (using the same parameters as described above) at the group level.

## Results

### Behavioural results

To ensure that participants were paying attention to and perceiving the directions of the RDKs, only participants with good performance (correlation between mean reported and presented directions > 0.9, see Materials and Methods for details) were included in the analysis (final sample: *r* = 0.98 +/- 0.02, mean +/- SD). In the localiser task, participants correctly detected dimming of the fixation dot with high accuracy (95.4% +/- 8.9%, mean +/- SD).

Participants’ perceptual reports of the moving dots’ direction was significantly biased towards the directions predicted by the auditory cues (F_1,92_ = 8.5, *p* = 0.0078; Figure 2). This indicates that, for identical visual stimuli, perception was partially determined by the predictive auditory cues. The cue-induced bias depended on the direction of the presented moving dots (F_4,92_ = 3.6, *p* = 0.0084), being weakest for close to horizontal (9°) and vertical (81°) directions, and stronger for directions closer to oblique (27° to 63°; see Figure 2).

### Expectations modulate sensory representations as early as 150ms post-stimulus

The first time-point at which motion direction could be decoded from the MEG signal evoked by task-irrelevant moving dot stimuli, in the localiser runs, was from 90 ms to 110 ms post-stimulus, peaking at 100 ms (*p*=0.018) (Figure 3). In order to probe modulations of early sensory signals by the predictive cues, motion direction decoding models were trained on participants’ individual peaks in this interval in the localiser runs, and applied to the main task (see Materials and Methods for details). Specifically, the decoded direction in trials where 27° was predicted was subtracted from the decoded direction in trials where 63° was predicted. This analysis revealed that across the whole group, the predictive auditory cues evoked a significant motion direction signal from 135 ms to 180 ms post-stimulus, peaking at 150 ms (peak difference=20.4°, *p*=0.027) (Figure 4A).

**Figure 4.**
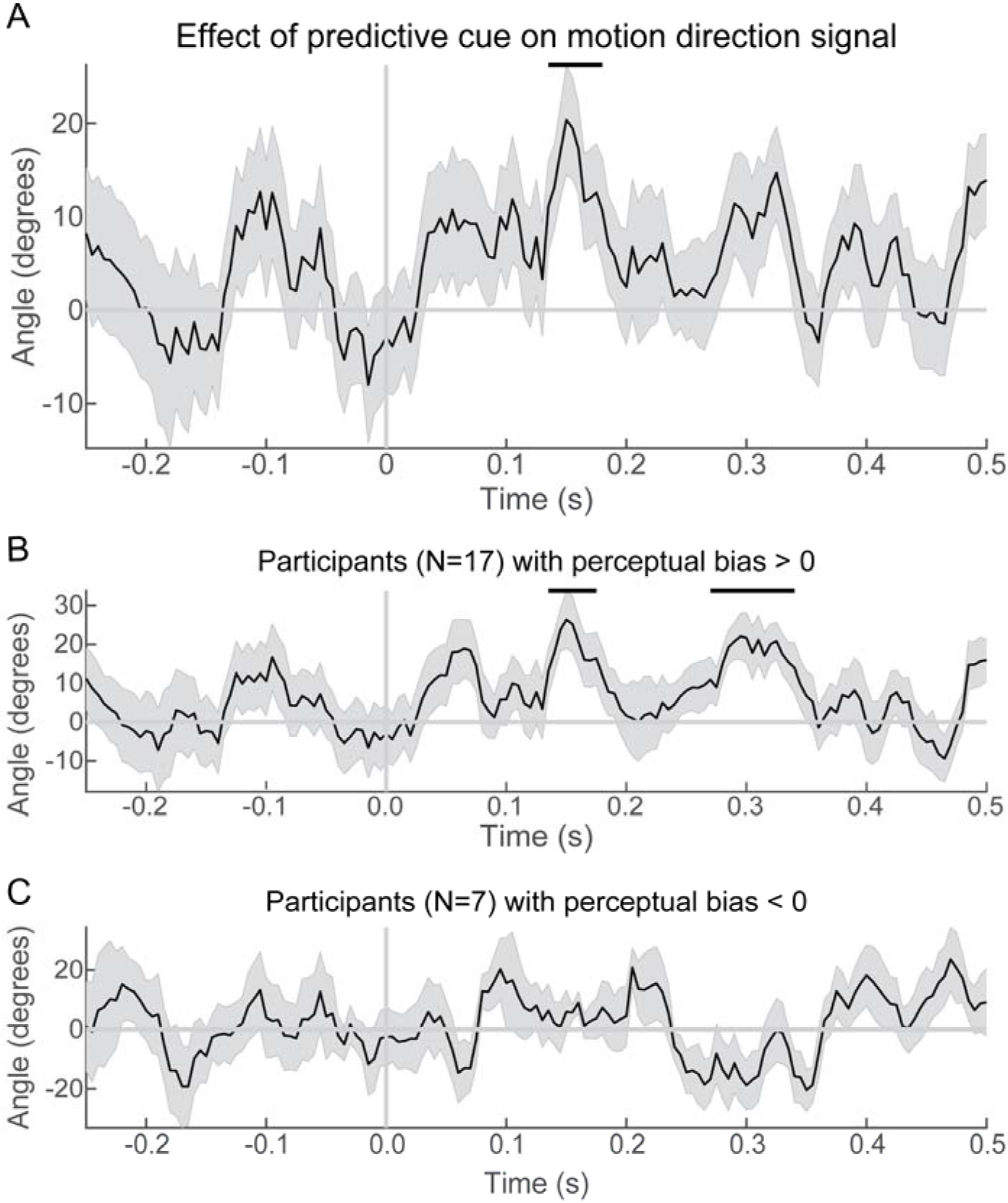
Expectation effects on decoded direction signals. **(A)** Linear subtraction of the decoded direction between the two cue conditions (i.e., 63° predicted – 27° predicted). **(B)** Cue effects on decoding restricted to participants who showed a perceptual bias towards the predicted directions. **(C)** Cue effects on decoding restricted to participants who did not show a perceptual bias towards the predicted directions. Shaded regions represent SEM; horizontal lines represent significant clusters (*p* < 0.05).

### Neural expectation signal related to perceptual bias

In order to relate this neural effect of the predictive cues to perception, we calculated it separately for participants who showed a positive perception bias (bias > 0°; N=17) and participants without a positive bias (bias <= 0°; N=7). In participants with an expectation-induced bias in perception, the predictive cues evoked a significant motion direction signal from 135 ms to 175 ms post-stimulus (peak = 26.3°, *p* = 0.044), as well as from 270 ms to 340 ms (peak = 22.0°, *p* = 0.0054) (Figure 4B). In the participants whose perception was not biased towards the predicted directions, there were no significant clusters in the MEG decoding signal (Figure 4C). In fact, participants with a perceptual bias towards the predicted directions displayed a significantly stronger prediction-evoked neural moving dot signal from 245 ms to 355 ms than participants without such a perceptual bias (*p* = 0.0005). Thus, individual variability in the neural signal evoked by the prediction cues was related to individual variability in the perceptual bias induced by these cues.

### Correlation between perceptual and neural representations emerges before stimulus onset

To further elucidate the relationship between neural sensory representations and perception, we correlated the decoded direction from the MEG data with the perceived direction on individual trials, controlling for the presented direction (i.e., through partial correlation, see Materials and Methods). This analysis aimed to investigate whether fluctuations in neural representations were related to fluctuations in subjective perception. For the participants whose perception was biased towards the predictive cues (N=17), decoded motion directions correlated significantly with perceived directions from −220 ms to 140 ms prestimulus (*p* = 0.0056) and 165 ms to 205 ms post-stimulus (*p* = 0.023) (Figure 5A). Thus, for these participants, the decoded neural signal was related to fluctuations in perception across trials. For the participants that did not display a perceptual bias towards the predictive cues (N=7), there were no significant clusters (Figure 5B). The correlation between decoded and perceived motion directions was significantly stronger for participants with a perceptual bias towards the predictive cues than for those without such a bias, from −10 to 35 ms (*p* = 0.041), and from 160 to 315 ms (*p* = 0.0009) post-stimulus. In other words, the relationship between neural and perceptual fluctuations was stronger in participants for whom the predictive cues induced an attractive perceptual bias.

**Figure 5.**
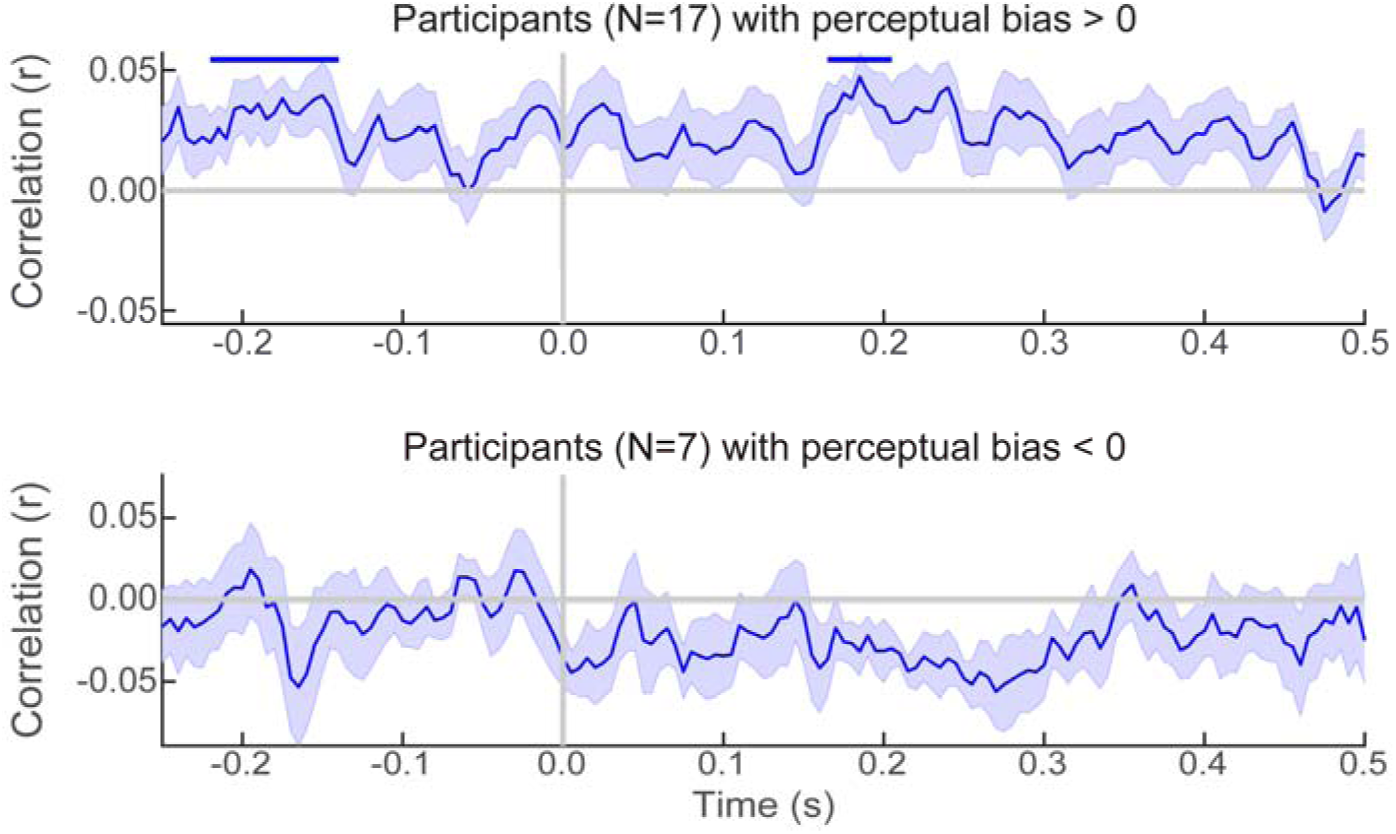
Single trial correlation between reported and decoded direction. Partial correlation between decoded and reported direction, controlling for presented direction **(A)** for participants with a perceptual bias towards the cues, **(B)** for participants without a perceptual bias towards the cues. Shaded regions represent SEM; horizontal lines represent significant clusters (*p* < 0.05).

### Implicit nature of the expectations

Participants were not informed of the meaning of the auditory cues, and all participants filled out a debriefing questionnaire to assess whether they were aware of the predictive relationships. Seven out of 24 participants indicated that they had noticed some relationship between the auditory tones and the direction of dots. Of these, only four correctly reported the true relationship, two reported the opposite relationship, and one was not able to report a specific relationship. The early neural effects of the predictive cues were still present when the four participants who reported being aware of the correct cue contingencies were excluded from the analysis (significant cluster from 125m s to 190 ms post-stimulus, *p* = 0.005), indicating that the expectation effects did not depend on the subjects being aware of the predictive relationships.

### Eye-movement control analysis

Eye-movements are known to be able to confound MEG decoding analyses (Mostert et al., 2018). In order to minimise the effects of eye movements, we estimated our decoding model based on independent localiser runs during which participants were performing a central-fixation task for which the directions of the moving dots were task-irrelevant (Mostert et al., 2018, Kok et al., 2017). Still, to investigate whether systematic eye movements during the localiser task could have affected our decoding results, we trained and tested the decoder on the vertical and horizontal gaze coordinates, as recorded by the eye tracker, rather than sensor amplitudes. Importantly, moving dot direction could not be decoded from the eye tracker signals in the first 150 ms post-stimulus (*r* < 0.1 for all time-points, no significant clusters at p < 0.05, also no single significant time-point between 0 and 150 ms, *p* < 0.05 uncorrected). This is the critical time window for our analyses, since the decoder was trained on MEG data from the localiser runs around 90 to 110 ms post-stimulus. As an additional stringent control, we performed the main analysis of interest on the eye-tracking data, that is, training the decoding models on the eye tracker signals in the localiser data on individual peaks between 90 and 110 ms, and applying them to the eye tracker data from the main task in order to reveal any effects of the predictive cues. As expected, since participants did not make systematic eye movements during this time window in the localiser, this analysis yielded no significant cue effects (no clusters at *p* < 0.05, also no single significant time-point between −250 and 500 ms, *p* < 0.05 uncorrected).

It is noteworthy that moving dot direction could be decoded from gaze position in the localiser at later time-points, namely from 170 ms to 360 ms post-stimulus (peak mean Pearson’s *r* = 0.460, *p* < 0.001). This indicates that participants moved their eyes systematically depending on the direction of the moving dots during this later time window in the localiser runs. Note however, as discussed above, that there were no significant points in the earlier time window in which we trained the decoder for the MEG data (90 ms to 110 ms), meaning that these later eye movements did not affect our analyses.

As a final control analysis, we trained the decoding models on the eye tracker data from the localiser runs on individual peaks between 170 and 200 ms post-stimulus (when localiser eye tracker decoding first became significant at the group level), and applied them to the eye tracker data from the main task. This analysis yielded no significant effects of the predictive cues on eye tracker signals (no clusters at *p* < 0.05, also no single significant time-point between −250 and 500 ms, *p* < 0.05 uncorrected).

In sum, while the moving dot stimuli were shown to affect gaze position at a later time window during the localiser runs, we found no evidence that systematic eye movements could explain the effects of interest here.

## Discussion

There has recently been much debate as to whether expectations can alter early sensory processing (De Lange et al., 2018; Summerfield and De Lange, 2014), or instead only modulate later decision-making processes (Rao et al., 2012; Rungratsameetaweemana et al., 2018). Here, we find that implicit prior expectations can modulate sensory representations at an early stage, in line with suggestions that expectations play a fundamental role in sensory processing (Summerfield and De Lange, 2014; Friston, 2005).

Perceptual reports of the direction of the moving dots were biased towards the direction predicted by the auditory cues. This is in line with previous studies (Kok et al., 2013; Chalk et al., 2010), as well as with theoretical work which casts perception as Bayesian inference, wherein the final percept is an integration of the prior expectations and perceptual input (Kersten et al., 2004; Knill and Richards, 1996). It is notable that expectations affected perception even though participants were not consciously aware of them (Kok et al. 2013; Chalk et al., 2010). However, while awareness of the predictive nature of the cues does not appear to be required in order to induce expectation effects in perception, awareness of the cues themselves may be (Meijs et al., 2018).

We found that the effect of the predictive cues depended on the presented direction: expectations affected perception more strongly when the presented direction was oblique than when it was near one of the cardinals (vertical or horizontal) (Figure 2). This may be because vertical and horizontal directions occur more frequently in natural environments, giving rise to ‘hyperpriors’: lifelong-learned expectations that these directions are likely to occur (Berkes et al., 2011; Girshick et al., 2011). Therefore at directions closer to the cardinal directions, the experimentally-induced priors may have interacted with hyperpriors, whereas at directions nearer 45° only the cue-related priors had an effect (Hu and Rahnev, 2019).

The primary motivation of this study was to establish whether expectations modulated the information content of early sensory signals. Predictive cues modulated the motion direction represented in the MEG signal as early as ∼150 ms post-stimulus, as revealed by a decoder trained on early (∼100 ms) MEG signals evoked by task-irrelevant moving dots in separate runs.

The relatively early time-point at which modulation occurs, and the sensory nature of the signal, are striking. The expectation signals were revealed by a decoder trained on task-irrelevant gratings, thus isolating sensory processes common between the localiser and main task (Kok et al., 2014; Kok et al., 2017). This, together with the fact that the decoders were trained on early (∼100 ms) post-stimulus time-points, suggests that these expectation signals reflect sensory processing, rather than being related to later decisional, attentional or motor processes (Mostert et al., 2015).

It should be noted that the latency at which expectation signals occurred (starting at 135 ms, peaking at 150 ms) is later than the first feedforward sweep of sensory information, which occurs within 50-80 ms (Alilović et al., 2019; Clark et al., 1994). It may therefore be that an initial feedforward sweep of sensory information is model-free, with expectations being integrated into the representation soon after (Alilović et al 2019; Marzecová et al., 2018). It is intriguing that many studies have found pre-stimulus effects of expectations (Alilović et al., 2019; Kok et al., 2017; Samaha et al., 2018) and attention (Myers et al., 2015); which do not seem to translate into subsequent modulations of the first feedforward sweep. The reason for this is not yet clear, but it has been suggested to be due to the fact that feedforward and feedback signals involve different neuronal populations (Alilović et al., 2019; Bastos et al., 2012; Kok et al., 2016). Alternatively, it may be that the lack of an earlier modulation by expectations was a result of the type of stimulus used: we used motion rather than static stimuli, meaning the accumulation of evidence takes inherently longer – indeed, within the localiser task, direction could not be decoded until 90 ms after stimulus onset, in contrast to studies using static stimuli which have shown significant decoding in the localiser as early as 40-60 ms post-stimulus (Kok et al., 2017; Mostert et al., 2015; Cichy et al., 2014). Furthermore, the decoding procedure used here was aimed at distinguishing several directions of motion within a ∼70° range (9° to 81°), rather than, for instance, decoding orthogonal orientations (Kok et al., 2017), which may have led to decreased signal-to-noise ratio, precluding successful decoding at earlier latencies.

In revealing expectation modulations of the information content of sensory signals at ∼150ms post-stimulus, our results accord with previous studies which report modulations of the amplitude of sensory signals by expectation at ∼100-150 ms (Samaha et al., 2018; Stojanoski and Niemeier, 2015; Bar et al., 2006; Todorovic et al. 2011; Todorovic and De Lange, 2012; Wacongne et al., 2011; Meyer and Olson, 2011; Aru et al., 2016). However, they conflict with experiments which reported no effects of expectation on early sensory processing (Rungratsameetaweemana et al., 2018; Rao et al., 2012). An important difference between studies may be the extent to which subjects actually form a perceptual expectation. For instance, in a perceptual decision-making study in macaques (Rao et al., 2012), the expectation cue predicted both which stimulus would appear, and what the correct task response was. Since the monkeys were strongly incentivised to perform the task accurately, the cue may have induced a response bias, rather than a perceptual bias. In the present study, to avoid strategic guessing or response bias, participants were not informed of the predictive relationship between the cues and the motion direction. In a recent study in humans using EEG that failed to find effects of expectation on sensory processing (Rungratsameetaweemana et al., 2018), the authors likely also predominantly manipulated task expectations, rather than perceptual expectations. That is, expectations pertained not so much to the statistics of the upcoming sensory inputs per se, but more so to which features of the inputs were likely to constitute a target. Therefore, this study more strongly manipulated task expectations (which features are likely to constitute a target) than perceptual expectations (which features are likely to appear on the screen).

A noteworthy aspect of our central finding is that the neural motion direction signal induced by the predictive cues peaked at around 20°, rising to 26° when considering only participants with a positive perceptual bias. This difference is an order of magnitude greater than the mean perceptual bias, being instead closer to the angle difference between the directions predicted by the two cues (63° vs 27°). This suggests that this early neural effect may reflect a reactivation of the predicted direction, rather than the integration of the predicted and presented directions. This is in line with the fact that this neural expectation effect peaks at ∼150 ms and then reduces, perhaps reflecting the integration of an expectation template (Kok et al., 2014; Kok et al., 2017) with incoming sensory evidence.

Intriguingly, for the participants who showed a perceptual bias towards the predictive cues (N=17), the neural expectation effect reappeared at ∼300 ms (Figure 4B). One possibility is that this reflects periodic activation of top-down expectations during recurrent feedforward and feedback message passing. Theoretical work which characterises perception as predictive processing postulates cycles of processing alternating between feedforward and feedback information, with the current hypothesis being tested and then iteratively revised until the hypothesis matches the incoming sensory information (Friston, 2005; Knill and Pouget, 2004; Bastos et al., 2012). Such recurrent message-passing has recently been shown to occur at a frequency of ∼11Hz during perception (Dijkstra et al., 2019).

Decoding analyses can reveal whether certain information is present in the neural signal, but not whether this information is used as a signal by the brain as part of the perceptual process, or is merely epiphenomenal. One way to address this inferential gap is to verify whether the decoded signal is related to behavioural variation (de-Wit et al., 2016). We found such a relationship both across participants and across trials. Across participants, neural expectation signals were stronger in participants with an expectation-induced perceptual bias. Across trials, for the participants with a perceptual bias towards the predicted directions, the correlation between decoded and perceived directions, regressing out the presented directions, was significant even before stimulus onset. Thus, the neural signal explained behavioural variation over and above that explained by the physical stimulus (Kok et al., 2013; St. John-Saaltink et al., 2016). This finding is in line with previous work suggesting the prestimulus state of the sensory cortex biases perception (Gandolfo and Downing, 2019; Kok et al., 2017; Sherman et al., 2016; Hesselmann et al, 2008; Pajani et al., 2015; Han and VanRullen, 2017).

In summary, our results demonstrate that expectations modulate the information content of sensory signals early on in the perceptual process. These findings are in line with predictive processing theories of perception that posit that expectations are a fundamental constituent of early sensory processing (Friston 2005; Lee and Mumford, 2003; Summerfield and De Lange, 2014; Keller and Mrsic-Flogel, 2018).

## Acknowledgements

We thank Daniel Bates for assistance with data collection. This work was supported by a Sir Henry Dale Fellowship jointly funded by the Wellcome Trust and the Royal Society (218535/Z/19/Z) to P.K. The Wellcome Centre for Human Neuroimaging is supported by core funding from the Wellcome Trust (203147/Z/16/Z).

## References

Alilović, J., Timmermans, B., Reteig, L. C., Van Gaal, S., & Slagter, H. A. (2019). No Evidence that Predictions and Attention Modulate the First Feedforward Sweep of Cortical Information Processing. Cerebral Cortex, 29(5), 2261–2278. https://doi.org/10.1093/cercor/bhz038

Alink, A., Schwiedrzik, C. M., Kohler, A., Singer, W., & Muckli, L. (2010). Stimulus Predictability Reduces Responses in Primary Visual Cortex. Journal of Neuroscience, 30(8), 2960–2966. https://doi.org/10.1523/JNEUROSCI.3730-10.2010

Aru, J., Rutiku, R., Wibral, M., Singer, W., & Melloni, L. (2016). Early effects of previous experience on conscious perception. Neuroscience of Consciousness, 2016(1). https://doi.org/10.1093/nc/niw004

Bang, J. W., & Rahnev, D. (2017). Stimulus expectation alters decision criterion but not sensory signal in perceptual decision making. Scientific Reports, 7(1), 17072. https://doi.org/10.1038/s41598-017-16885-2

Bar, M., Kassam, K. S., Ghuman, A. S., Boshyan, J., Schmid, A. M., Dale, A. M., Hämäläinen, M. S., Marinkovic, K., Schacter, D. L., Rosen, B. R., & Halgren, E. (2006). Top-down facilitation of visual recognition. Proceedings of the National Academy of Sciences of the United States of America, 103(2), 449–454. https://doi.org/10.1073/pnas.0507062103

Bastos, A. M., Usrey, W. M., Adams, R. A., Mangun, G. R., Fries, P., & Friston, K. J. (2012). Canonical microcircuits for predictive coding. Neuron, 76(4), 695–711. https://doi.org/10.1016/j.neuron.2012.10.038

Blankertz, B., Lemm, S., Treder, M., Haufe, S., & Müller, K.-R. (2011). Single-trial analysis and classification of ERP components—A tutorial. NeuroImage, 56(2), 814–825. https://doi.org/10.1016/j.neuroimage.2010.06.048

Brouwer, G. J., & Heeger, D. J. (2009). Decoding and Reconstructing Color from Responses in Human Visual Cortex. Journal of Neuroscience, 29(44), 13992–14003. https://doi.org/10.1523/JNEUROSCI.3577-09.2009

Brouwer, Gijs Joost, & Heeger, D. J. (2011). Cross-orientation suppression in human visual cortex. Journal of Neurophysiology, 106(5), 2108–2119. https://doi.org/10.1152/jn.00540.2011

Chalk, M., Seitz, A. R., & Seriès, P. (2010). Rapidly learned stimulus expectations alter perception of motion. Journal of Vision, 10(8), 2–2. https://doi.org/10.1167/10.8.2

Chaumon, M., Drouet, V., & Tallon-Baudry, C. (2008). Unconscious associative memory affects visual processing before 100 ms. Journal of Vision, 8(3), 10–10. https://doi.org/10.1167/8.3.10

Cichy, R. M., Pantazis, D., & Oliva, A. (2014). Resolving human object recognition in space and time. Nature Neuroscience, 17(3), 455–462. https://doi.org/10.1038/nn.3635

Clark, V. P., Fan, S., & Hillyard, S. A. (1994). Identification of early visual evoked potential generators by retinotopic and topographic analyses. Human Brain Mapping, 2(3), 170–187. https://doi.org/10.1002/hbm.460020306

De Lange, F. P., Heilbron, M., & Kok, P. (2018). How Do Expectations Shape Perception? Trends in Cognitive Sciences, 22(9), 764–779.

de-Wit, L., Alexander, D., Ekroll, V., & Wagemans, J. (2016). Is neuroimaging measuring information in the brain? Psychonomic Bulletin & Review, 23(5), 1415–1428. https://doi.org/10.3758/s13423-016-1002-0

Den Ouden, H. E. M., Friston, K. J., Daw, N. D., McIntosh, A. R., & Stephan, K. E. (2009). A Dual Role for Prediction Error in Associative Learning. Cerebral Cortex, 19(5), 1175–1185. https://doi.org/10.1093/cercor/bhn161

Dijkstra, N., Ambrogioni, L., & van Gerven, M. A. J. (2019). Neural dynamics of perceptual inference and its reversal during imagery [Preprint]. bioRxiv. https://doi.org/10.1101/781294

Fisher, R. A. (1915). Frequency Distribution of the Values of the Correlation Coefficient in Samples from an Indefinitely Large Population. Biometrika, 10(4), 507–521. https://doi.org/10.2307/2331838

Friston, K. (2005). A theory of cortical responses. Philosophical Transactions of the Royal Society B: Biological Sciences, 360(1456), 815–836. https://doi.org/10.1098/rstb.2005.1622

Gamond, L., George, N., Lemaréchal, J.-D., Hugueville, L., Adam, C., & Tallon-Baudry, C. (2011). Early influence of prior experience on face perception. NeuroImage, 54(2), 1415–1426. https://doi.org/10.1016/j.neuroimage.2010.08.081

Gandolfo, M., & Downing, P. E. (2019). Causal Evidence for Expression of Perceptual Expectations in Category-Selective Extrastriate Regions. Current Biology, 29(15), 2496-2500.e3. https://doi.org/10.1016/j.cub.2019.06.024

Garcia, J. O., Srinivasan, R., & Serences, J. T. (2013). Near-real-time feature-selective modulations in human cortex. Current Biology, 23(6), 515–522. https://doi.org/10.1016/j.cub.2013.02.013

Girshick, A. R., Landy, M. S., & Simoncelli, E. P. (2011). Cardinal rules: Visual orientation perception reflects knowledge of environmental statistics. Nature Neuroscience, 14(7), 926–932. https://doi.org/10.1038/nn.2831

Gold, J. I., & Shadlen, M. N. (2007). The Neural Basis of Decision Making. Annual Review of Neuroscience, 30(1), 535–574. https://doi.org/10.1146/annurev.neuro.29.051605.113038

Han, B., & VanRullen, R. (2017). The rhythms of predictive coding? Pre-stimulus phase modulates the influence of shape perception on luminance judgments. Scientific Reports, 7(1), 1–10. https://doi.org/10.1038/srep43573

Harrison, S. A., & Tong, F. (2009). Decoding reveals the contents of visual working memory in early visual areas. Nature, 458(7238), 632–635. https://doi.org/10.1038/nature07832

Heekeren, H. R., Marrett, S., Bandettini, P. A., & Ungerleider, L. G. (2004). A general mechanism for perceptual decision-making in the human brain. Nature, 431(7010), 859–862. https://doi.org/10.1038/nature02966

Helmholtz, H. von. (2013). Treatise on Physiological Optics. Courier Corporation.

Hesselmann, G., Kell, C. A., Eger, E., & Kleinschmidt, A. (2008). Spontaneous local variations in ongoing neural activity bias perceptual decisions. Proceedings of the National Academy of Sciences of the United States of America, 105(31), 10984–10989. https://doi.org/10.1073/pnas.0712043105

Hsu, Y.-F., Le Bars, S., Hamalainen, J. A., & Waszak, F. (2015). Distinctive Representation of Mispredicted and Unpredicted Prediction Errors in Human Electroencephalography. Journal of Neuroscience, 35(43), 14653–14660. https://doi.org/10.1523/JNEUROSCI.2204-15.2015

Hu, M., & Rahnev, D. (2019). Predictive cues reduce but do not eliminate intrinsic response bias. Cognition, 192, 104004. https://doi.org/10.1016/j.cognition.2019.06.016

Jabar, S. B., Filipowicz, A., & Anderson, B. (2017). Tuned by experience: How orientation probability modulates early perceptual processing. Vision Research, 138, 86–96. https://doi.org/10.1016/j.visres.2017.07.008

Keller, G. B., & Mrsic-Flogel, T. D. (2018). Predictive Processing: A Canonical Cortical Computation. Neuron, 100(2), 424–435. https://doi.org/10.1016/j.neuron.2018.10.003

Kersten, D., Mamassian, P., & Yuille, A. (2004). Object Perception as Bayesian Inference. Annual Review of Psychology, 55(1), 271–304. https://doi.org/10.1146/annurev.psych.55.090902.142005

Kersten, D., & Yuille, A. (2003). Bayesian models of object perception. Current Opinion in Neurobiology, 13(2), 150–158. https://doi.org/10.1016/s0959-4388(03)00042-4

Knill, D. C., & Pouget, A. (2004). The Bayesian brain: The role of uncertainty in neural coding and computation. Trends in Neurosciences, 27(12), 712–719. https://doi.org/10.1016/j.tins.2004.10.007

Knill, D. C., & Richards, W. (1996). Perception as Bayesian Inference. Cambridge University Press.

Kok, P., Bains, L. J., van Mourik, T., Norris, D. G., & de Lange, F. P. (2016). Selective Activation of the Deep Layers of the Human Primary Visual Cortex by Top-Down Feedback. Current Biology, 26(3), 371–376. https://doi.org/10.1016/j.cub.2015.12.038

Kok, P., Brouwer, G. J., van Gerven, M. A. J., & de Lange, F. P. (2013). Prior Expectations Bias Sensory Representations in Visual Cortex. The Journal of Neuroscience, 33(41), 16275–16284. https://doi.org/10.1523/JNEUROSCI.0742-13.2013

Kok, P., Failing, M. F., & de Lange, F. P. (2014). Prior Expectations Evoke Stimulus Templates in the Primary Visual Cortex. Journal of Cognitive Neuroscience, 26(7), 1546–1554. https://doi.org/10.1162/jocn_a_00562

Kok, P., Jehee, J. F. M., & de Lange, F. P. (2012). Less Is More: Expectation Sharpens Representations in the Primary Visual Cortex. Neuron, 75(2), 265–270. https://doi.org/10.1016/j.neuron.2012.04.034

Kok, P., Mostert, P., & de Lange, F. P. (2017). Prior expectations induce prestimulus sensory templates. Proceedings of the National Academy of Sciences of the United States of America, 114(39), 10473–10478. https://doi.org/10.1073/pnas.1705652114

Lee, T. S., & Mumford, D. (2003). Hierarchical Bayesian inference in the visual cortex. Journal of the Optical Society of America A, 20(7), 1434. https://doi.org/10.1364/JOSAA.20.001434

Maris, E., & Oostenveld, R. (2007). Nonparametric statistical testing of EEG- and MEG-data. Journal of Neuroscience Methods, 164(1), 177–190. https://doi.org/10.1016/j.jneumeth.2007.03.024

Marzecová, A., Schettino, A., Widmann, A., SanMiguel, I., Kotz, S. A., & Schröger, E. (2018). Attentional gain is modulated by probabilistic feature expectations in a spatial cueing task: ERP evidence. Scientific Reports, 8(1), 54. https://doi.org/10.1038/s41598-017-18347-1

Meijs, E. L., Slagter, H. A., de Lange, F. P., & van Gaal, S. (2018). Dynamic Interactions between Top– Down Expectations and Conscious Awareness. The Journal of Neuroscience, 38(9), 2318–2327. https://doi.org/10.1523/JNEUROSCI.1952-17.2017

Meyer, T., & Olson, C. R. (2011). Statistical learning of visual transitions in monkey inferotemporal cortex. Proceedings of the National Academy of Sciences of the United States of America, 108(48), 19401–19406. https://doi.org/10.1073/pnas.1112895108

Mostert, P., Albers, A. M., Brinkman, L., Todorova, L., Kok, P., & de Lange, F. P. (2018). Eye Movement-Related Confounds in Neural Decoding of Visual Working Memory Representations. eNeuro, 5(4). https://doi.org/10.1523/ENEURO.0401-17.2018

Mostert, P., Kok, P., & de Lange, F. P. (2016). Dissociating sensory from decision processes in human perceptual decision making. Scientific Reports, 5(1), 18253. https://doi.org/10.1038/srep18253

Myers, N. E., Rohenkohl, G., Wyart, V., Woolrich, M. W., Nobre, A. C., & Stokes, M. G. (2015). Testing sensory evidence against mnemonic templates. eLife, 4, e09000. https://doi.org/10.7554/eLife.09000

Oostenveld, R., Fries, P., Maris, E., & Schoffelen, J.-M. (2011). FieldTrip: Open Source Software for Advanced Analysis of MEG, EEG, and Invasive Electrophysiological Data. Computational Intelligence and Neuroscience, 2011. https://doi.org/10.1155/2011/156869

Pajani, A., Kok, P., Kouider, S., & de Lange, F. P. (2015). Spontaneous Activity Patterns in Primary Visual Cortex Predispose to Visual Hallucinations. The Journal of Neuroscience, 35(37), 12947–12953. https://doi.org/10.1523/JNEUROSCI.1520-15.2015

Rao, R. P. N., & Ballard, D. H. (1999). Predictive coding in the visual cortex: A functional interpretation of some extra-classical receptive-field effects. Nature Neuroscience, 2(1), 79–87. https://doi.org/10.1038/4580

Rao, V., DeAngelis, G. C., & Snyder, L. H. (2012). Neural Correlates of Prior Expectations of Motion in the Lateral Intraparietal and Middle Temporal Areas. The Journal of Neuroscience, 32(29), 10063–10074. https://doi.org/10.1523/JNEUROSCI.5948-11.2012

Rungratsameetaweemana, N., Itthipuripat, S., Salazar, A., & Serences, J. T. (2018). Expectations Do Not Alter Early Sensory Processing during Perceptual Decision-Making. The Journal of Neuroscience, 38(24), 5632–5648. https://doi.org/10.1523/JNEUROSCI.3638-17.2018

Samaha, J., Boutonnet, B., Postle, B. R., & Lupyan, G. (2018). Effects of meaningfulness on perception: Alpha-band oscillations carry perceptual expectations and influence early visual responses. Scientific Reports, 8. https://doi.org/10.1038/s41598-018-25093-5

Sherman, M. T., Kanai, R., Seth, A. K., & VanRullen, R. (2016). Rhythmic Influence of Top–Down Perceptual Priors in the Phase of Prestimulus Occipital Alpha Oscillations. Journal of Cognitive Neuroscience, 28(9), 1318–1330. https://doi.org/10.1162/jocn_a_00973

St John-Saaltink, E., Kok, P., Lau, H. C., & de Lange, F. P. (2016). Serial Dependence in Perceptual Decisions Is Reflected in Activity Patterns in Primary Visual Cortex. The Journal of Neuroscience, 36(23), 6186–6192. https://doi.org/10.1523/JNEUROSCI.4390-15.2016

Stojanoski, B. B., & Niemeier, M. (2015). Colour expectations during object perception are associated with early and late modulations of electrophysiological activity. Experimental Brain Research, 233(10), 2925–2934. https://doi.org/10.1007/s00221-015-4362-1

Summerfield, C., & de Lange, F. P. (2014). Expectation in perceptual decision making: Neural and computational mechanisms. Nature Reviews Neuroscience, 15(11), 745–756. https://doi.org/10.1038/nrn3838

Todorovic, A., & de Lange, F. P. (2012). Repetition suppression and expectation suppression are dissociable in time in early auditory evoked fields. The Journal of Neuroscience, 32(39), 13389–13395. https://doi.org/10.1523/JNEUROSCI.2227-12.2012

Todorovic, A., van Ede, F., Maris, E., & de Lange, F. P. (2011). Prior expectation mediates neural adaptation to repeated sounds in the auditory cortex: An MEG study. The Journal of Neuroscience, 31(25), 9118–9123. https://doi.org/10.1523/JNEUROSCI.1425-11.2011

Wacongne, C., Labyt, E., Wassenhove, V. van, Bekinschtein, T., Naccache, L., & Dehaene, S. (2011). Evidence for a hierarchy of predictions and prediction errors in human cortex. Proceedings of the National Academy of Sciences of the United States of America, 108(51), 20754–20759. https://doi.org/10.1073/pnas.1117807108

Wyart, V., Nobre, A. C., & Summerfield, C. (2012). Dissociable prior influences of signal probability and relevance on visual contrast sensitivity. Proceedings of the National Academy of Sciences of the United States of America, 109(9), 3593–3598. https://doi.org/10.1073/pnas.1120118109

Xu, Y. (2018). Sensory Cortex Is Nonessential in Working Memory Storage. Trends in Cognitive Sciences, 22(3), 192–193. https://doi.org/10.1016/j.tics.2017.12.008

Zhang, X., Yan, W., Wang, W., Fan, H., Hou, R., Chen, Y., Chen, Z., Ge, C., Duan, S., Compte, A., & Li, C. T. (2019). Active information maintenance in working memory by a sensory cortex. eLife, 8, e43191. https://doi.org/10.7554/eLife.43191

